# An important role for triglyceride in regulating spermatogenesis

**DOI:** 10.1101/2022.12.16.520841

**Authors:** Charlotte F. Chao, Yanina-Yasmin Pesch, Huaxu Yu, Chenjingyi Wang, Maria J. Aristizabal, Tao Huan, Guy Tanentzapf, Elizabeth J. Rideout

## Abstract

*Drosophila* is a powerful model to study how lipids affect spermatogenesis. Yet, the contribution of neutral lipids, a major lipid group which resides in organelles called lipid droplets (LD), to sperm development is largely unknown. Emerging evidence suggests LD are present in the testis and that loss of neutral lipid- and LD-associated genes causes subfertility; however, key regulators of testis neutral lipids and LD remain unclear. Here, we show LD are present in early-stage somatic and germline cells within the *Drosophila* testis. We identified a role for triglyceride lipase *brummer* (*bmm*) in regulating testis LD, and found that whole-body loss of *bmm* leads to defects in sperm development. Importantly, these represent cell-autonomous roles for *bmm* in regulating testis LD and spermatogenesis. Because lipidomic analysis of *bmm* mutants revealed excess triglyceride accumulation, and spermatogenic defects in *bmm* mutants were rescued by genetically blocking triglyceride synthesis, our data suggest that *bmm*-mediated regulation of triglyceride influences sperm development. This identifies triglyceride as an important neutral lipid that contributes to *Drosophila* sperm development, and reveals a key role for *bmm* in regulating testis triglyceride levels during spermatogenesis.

## INTRODUCTION

Lipids play an essential role in regulating spermatogenesis across animals [1–4]. Studies in *Drosophila* have illuminated key roles for multiple lipid species in regulating sperm development [5–7]. For example, phosphatidylinositol and its phosphorylated derivatives participate in diverse aspects of *Drosophila* spermatogenesis including meiotic cytokinesis [1,8–11], somatic cell differentiation [12], germline and somatic cell polarity maintenance [13–16], and germline stem cell (GSC) maintenance and proliferation [17]. Membrane lipids also influence sperm development [18,19], whereas fatty acids play a role in processes such as meiotic cytokinesis [20] and sperm individualization [21,22]. While these studies suggest key roles for membrane lipids and fatty acids during *Drosophila* spermatogenesis, some of which are conserved in mammals [23–25], much less is known about how neutral lipids contribute to spermatogenesis.

Neutral lipids are a major lipid group that includes triglyceride and cholesterol ester, and reside within specialized organelles called lipid droplets (LD) [26]. LD are found in diverse cell types (*e.g*. adipocytes, muscle, liver, glia, neurons) [27,28,26], and play key roles in maintaining cellular lipid homeostasis. In nongonadal cell types, correct regulation of LD contributes to cellular energy production [29–31], sequestration and redistribution of lipid precursors [32–36], and regulation of lipid toxicity [37–39]. The importance of LD to normal cellular function in nongonadal cell types is shown by the fact that dysregulation of LD causes defects in cell differentiation, survival, and energy production [26,37,40,41]. In the testis, much less is known about the regulation and function of neutral lipids and LD, and how this regulation affects sperm development.

Multiple lines of evidence suggest a potential role for neutral lipids and LD during spermatogenesis. First, genes that encode proteins associated with neutral lipid metabolism and LD are expressed in the testis across multiple species [42–44]. Second, testis LD have been identified in mammals and flies under both normal physiological conditions [27,44–48] and after mitochondrial stress [49]. Third, loss of genes associated with neutral lipid metabolism and LD cause subfertility phenotypes in both flies and mammals [27,50–52]. While studies suggest that mammalian testis LD contribute to steroidogenesis [53,54], the spatial, temporal, and cell-type specific requirements for neutral lipids and LD in the testis have not been explored in detail in any animal. It remains similarly unclear which genes are responsible for regulating neutral lipids and LD during spermatogenesis.

To address these knowledge gaps, we used *Drosophila* to investigate the regulation and function of neutral lipids and LD during sperm development. Our detailed analysis of spermatogenesis under normal physiological conditions revealed the presence of LD in early-stage somatic and germline cells in the testis. We identified triglyceride lipase *brummer* (*bmm*) as a regulator of testis LD, and showed that this represents a cell-autonomous role for *bmm* in the germline. Importantly, we found that *bmm*-mediated regulation of testis LD was significant for spermatogenesis, as both whole-body and cell-autonomous loss of *bmm* caused defects in sperm development. Given that our lipidomic analysis revealed an excess accumulation of triglyceride in animals lacking *bmm*, and that genetically blocking triglyceride synthesis rescued many spermatogenic defects associated with *bmm* loss, our data suggests that *bmm*-mediated regulation of triglyceride is important for normal *Drosophila* sperm development. This reveals previously unrecognized roles for neutral lipids such as triglyceride in regulating spermatogenesis, and for *bmm* in regulating sperm development under normal physiological conditions. Together, these findings advance knowledge of the regulation and function of neutral lipids during spermatogenesis.

## RESULTS

### Lipid droplets are present in early-stage somatic and germline cells

We previously reported the presence of small LD (<1 μm) at the apical tip of the testis [27], a finding we reproduced in *w^1118^* males using neutral lipid stain BODIPY (Figure 1A). These LD were present in the region that contains stem cells, early-stage somatic cells, and germline cells (Figure 1A-A’, arrows). LD were also present in the hub, an organizing center and stem cell niche in the *Drosophila* testis (Figure 1A’’-A’’’, arrows) [55], but largely absent within the area occupied by spermatocytes (Figure 1A and A’, arrowheads). This LD distribution was reproduced in two independent genetic backgrounds and at two additional ages (Figure 1B, 1C). While LD may contain multiple neutral lipid species [56], cholesterol-binding fluorescent polyene antibiotic filipin III did not detect cholesterol within testis LD (Figure S1A), suggesting triglyceride is the main neutral lipid in *Drosophila* testis LD.

**Figure 1.**
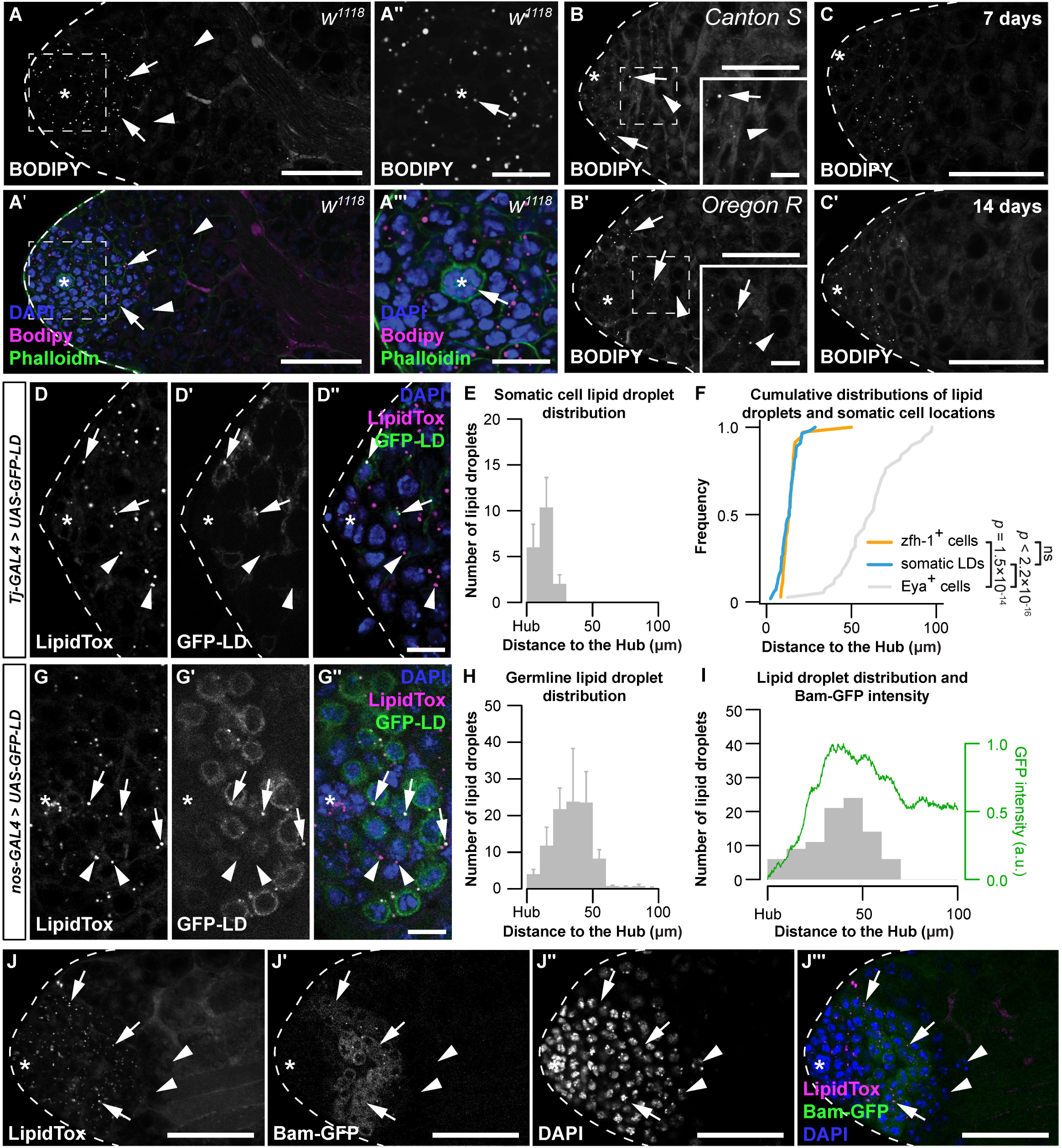
Lipid droplets are present in early-stage somatic and germ cells. (A) Testis lipid droplets (LD) in *w^1118^* animals visualized with neutral lipid dye BODIPY. (A,A’) Scale bar=50 μm; (A’’,A’’’) scale bar=15 μm. Asterisk indicates hub in all images. Arrows point to LD; arrowheads point to spermatocytes in A,B. Spermatocytes were identified as described in methods section. (B) Testis LD visualized with BODIPY in newly-eclosed males from two wild-type genotypes. Scale bars: main image=50 μm; inset image=10 μm. (C) Testis LD from *w^1118^* animals at different times post-eclosion. Scale bars=50 μm. (D) Testis LD visualized with LipidTox Red in animals with somatic cell overexpression of GFP-LD (*Tj-GAL4*>*UAS-GFP-LD*). GFP- and LipidTox Red-positive punctae are somatic LD (D–D’’ arrows); LipidTox punctae without GFP indicate germline LD (D–D’’ arrowheads). Scale bars=10 μm. (E) Histogram showing the spatial distribution of somatic cell LD; error bars represent standard error of the mean (SEM). (F) Cumulative frequency distributions of somatic LD (blue line, data reproduced from E), zfh-1-positive somatic cells (zfh-1^+^ cells, orange line), and Eya-positive somatic cells (Eya^+^ cells, grey line). (G) Testis LD visualized with LipidTox Red in males with germline overexpression of GFP-LD (*nos-GAL4*>*UAS-GFP-LD*). GFP- and LipidTox Red-positive punctae indicate germline LD (arrows); LipidTox punctae without GFP indicate non-germline LD (arrowheads). Scale bars=10 μm. (H) Histogram representing the spatial distribution of LD within the germline; error bars represent SEM. (I) Histogram representing the spatial distribution of LD and GFP fluorescence (green line) (arbitrary units, a.u.) in a representative testis of a *bam-GFP* animal (panel J). (J) Testis LD in a *bam-GFP* animal; arrows point to LD and arrowheads point to spermatocytes. Scale bar=50 μm. See also Supplemental Figure 1.

*Drosophila* spermatogenesis requires the co-development and differentiation of the germline and the somatic lineages [57]. To identify LD in each lineage, we used the GAL4/UAS system to overexpress a GFP transgene fused to the LD-targeting motif of motor protein *Klarsicht* (*UAS*-*GFP-LD*) [58]. Somatic overexpression of *UAS-GFP-LD* using *Traffic jam (Tj)-GAL4* revealed that the majority of somatic LD in 0-day-old males were located <30 μm from the hub (Figure 1D, 1E). Because the somatic LD distribution coincided with a marker for somatic stem cells and their immediate daughter cells (Zinc finger homeodomain 1, Zfh-1) (Figure 1F; two-sample Kolmogorov-Smirnov test) [59], but not with a marker for late somatic cells (Eyes absent, Eya) [12,60], our data suggest LD are present in early somatic cells. Germline overexpression of *UAS-GFP-LD* using *nanos (nos)-GAL4* demonstrated the presence of LD within germline cells near the apical tip of the testis in 0-day-old males (Figure 1G, 1H). Specifically, the disappearance of germline LD coincided with peak expression of a GFP reporter that reflects the expression of Bag-of-marbles (Bam) protein in the testis (Bam-GFP) [61] (Figure 1I, 1J). Because peak Bam expression signals the last round of transient amplifying mitotic cell cycle prior to the germline’s transition into the meiotic cell cycle [62–64], our data suggests that germline LD, like somatic LD, are present in cells at early stages of development.

### *brummer* plays a cell-autonomous role in regulating testis lipid droplets

*Adipose triglyceride lipase* (*ATGL*) is a critical regulator of neutral lipid metabolism and LD [65–74]. Loss of *ATGL* in many cell types triggers LD accumulation, and *ATGL* overexpression decreases LD number [30,67,68,71,73,75,76]. Given that the *Drosophila ATGL* homolog *brummer* (*bmm*) regulates testis LD induced by mitochondrial stress [49], we explored whether *bmm* regulates testis LD under normal physiological conditions. We first examined *bmm* expression in the testis by isolating this organ from flies carrying a *bmm* promoter-driven *GFP* transgene (*bmm-GFP*) that recapitulates many aspects of *bmm* mRNA regulation [77]. GFP expression was present in the germline of *bmm-GFP* testes, and we found germline GFP levels were higher in spermatocytes than at earlier stages of sperm development (Figure 2A, 2B; one-way ANOVA with Tukey multiple comparison test). In further support of this observation, we analyzed a publicly available single-cell RNA sequencing dataset from the male reproductive organ [78]. Using pseudotime analysis, we arranged the germline (Figure S2A) and somatic cells (Figure S2B) based on their annotated developmental trajectory. The expression pattern of *bmm* in the germline matched our observation with the *bmm-GFP* reporter (Figure S2C). While levels of the *bmm-GFP* reporter were lower in somatic cells, single-cell RNA sequencing data identified *bmm* expression in the somatic lineage that was higher in cells at later stages of development (Figure S2D). Additional neutral lipid- and lipid droplet-associated genes such as *lipid storage droplet*-*2*, *Seipin*, *Lipin*, and *midway* also showed differential regulation during differentiation in the testis (Figure S2C, S2D). Combined with our data on the location of testis LD, these gene expression data suggest that *bmm* upregulation in both somatic and germline cells during differentiation corresponds to the downregulation of testis LD. Supporting this, germline GFP levels were negatively correlated with testis LD in *bmm-GFP* flies (Figure 2A, 2C), suggesting regions with higher *bmm* expression had fewer LD.

**Figure 2.**
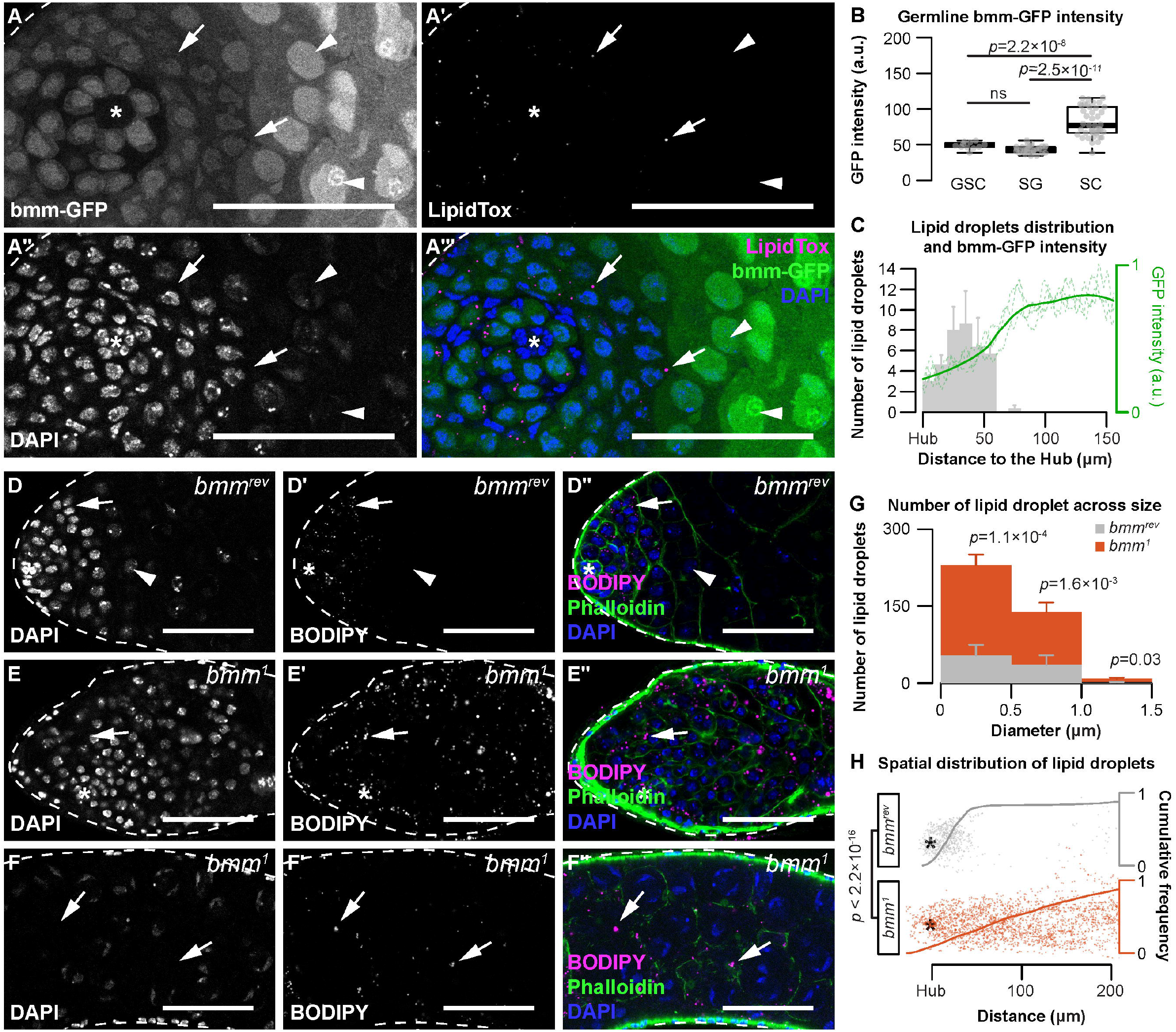
*brummer* regulates lipid droplets in both germline and somatic cells of the testis. (A-A’’’’) Testis lipid droplets (LD) indicated by LipidTox Red in *bmm-GFP* animals. Arrows point to LD in all images. Arrowheads point to spermatocytes. Scale bars=50 μm. Asterisks indicate the hub in all images. (B) Quantification of nuclear GFP intensity in testes isolated from *bmm-GFP* animals (n=3). Germline stem cell (GSC), spermatogonia (SG), spermatocyte (SC). (C) Spatial distribution of LD (grey histogram) and GFP expression (green line) in testes from *bmm-GFP* animals as a function of distance from the hub (n=3). (D,E) LD near the apical region of the testis in *bmm^rev^* (D) or *bmm^1^* (E) animals. (F) LD further away from the apical tip in *bmm^1^* animals. (D–F) Scale bars=50 μm. (G) Histogram representing testis LD size distribution in *bmm^rev^* (grey) and *bmm^1^* (orange). (H) Apical tip of the testes is at the left of the graph; individual dots represent a single LD and its relative position to the hub marked by an asterisk. Cumulative frequency distribution of the distance between LDs and the apical tip of the testes are drawn as solid lines. See also Supplemental Figure 2 and 3.

To test whether *bmm* regulates testis LD, we compared LD in testes from 0-day-old males carrying a loss-of-function mutation in *bmm* (*bmm^1^*) to control male testes (*bmm^rev^)* [67]. *bmm^1^* males had significantly more LD across all LD sizes compared with control males at the apical tip of the testis (Figure 2D–2G; Welch two-sample t-test with Bonferroni correction) and showed a significantly expanded LD distribution (Figure 2D– 2F, 2H; two-sample Kolmogorov-Smirnov test). This suggests *bmm* normally restricts LD to the apical tip of the testis, an observation we confirmed in both somatic and germline lineages using lineage-specific expression of GFP-LD (Figure S3A–S3D). Importantly, after inducing homozygous *bmm^rev^* or *bmm^1^* clones in the testes using the *FLP*-*FRT* system (Figure 3A, 3B) [79], we found *bmm^1^* spermatocyte clones had significantly more LD at 3 days post-clone induction (Figure 3C; Welch two-sample t-test), a stage at which LD were absent from *bmm^rev^* clones. Because we observed no significant effect of cell-autonomous *bmm* loss on LD at any other stage of germline development (Figure 3C), this suggests *bmm* function is not required to regulate LD at early stages of germ cell development. Instead, our data suggest *bmm* plays a role in regulating LD at the spermatogonia-spermatocyte transition. While we were unable to assess LD in *bmm^1^* somatic clones, our data reveals a previously unrecognized cell-autonomous role for *bmm* as a regulator of LD in germline cells.

**Figure 3.**
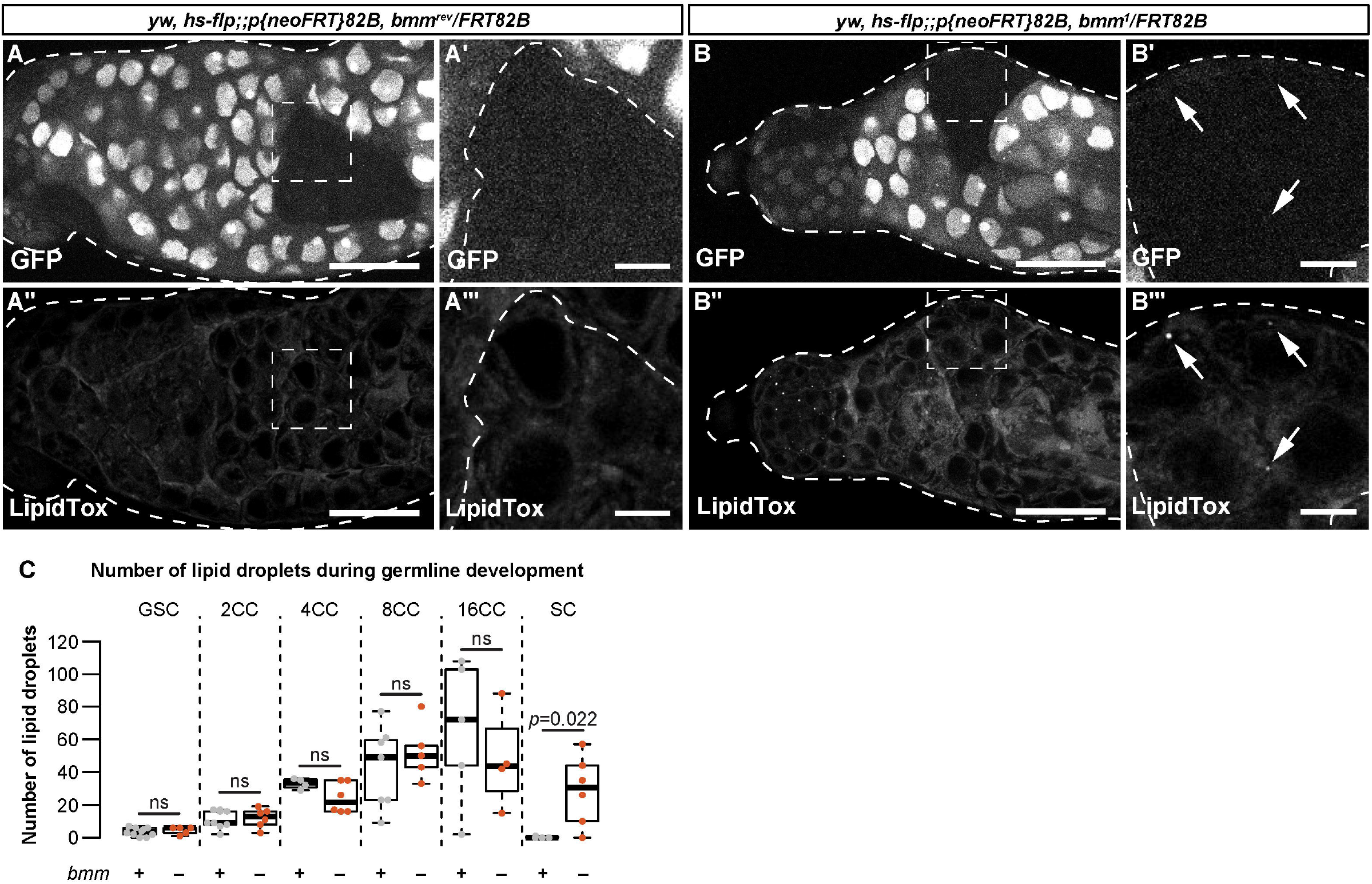
*bmm* regulates germline lipid droplets in a cell-autonomous manner. (A–B) Single confocal slices through a representative testis isolated from an individual carrying clones induced using the FLP-FRT system at 3 days post-clone induction. Clones are homozygous for an allele that encodes a functional *bmm* protein product (*bmm^rev^*; A–A’’’) or a loss-of-function *bmm* allele (*bmm^1^,* B–B’’’). GFP negative areas mark homozygous clones in panel A and B; the boxed areas in A, A’’ and B, B’’ are shown in A’, A’’’ and B’, B’’’, respectively. In homozygous *bmm^rev^* spermatocyte clones we detected no LD using neutral lipid dye LipidTox (A’’ and A’’’). In contrast, spermatocyte clones homozygous for *bmm^1^* have detectable LD (B’’ and B’’’, arrows). Scale bars=50 μm in A,A’’ and B,B’’; scale bars=10 μm in A’,A’’’ and B’,B’’’. (C) Number of testis LD in *bmm^rev^* (grey) or *bmm^1^* (orange) in FLP-FRT clones 3 days post-clone induction; dots represent measurements from a single clone. The number of cells in each cyst (CC) counted is indicated. There were significantly more LD in *bmm^1^* spermatocyte (SC) clones (*p* = 0.026; Welch two-sample t-test) but not at other stages of development.

### *brummer* plays a cell-autonomous role in regulating germline development

To determine the physiological significance of *bmm-*mediated regulation of testis LD, we investigated testis and sperm development in males without *bmm* function. In 0-day-old *bmm^1^* males reared at 25°C, testis size was significantly smaller than in age-matched *bmm^rev^* controls (Figure 4A, 4B; Welch two-sample t-test), and the number of spermatid bundles was significantly lower (Figure 4C; Kruskal-Wallis rank sum test). When the animals were reared at 29°C, a temperature that exacerbates spermatogenesis defects associated with changes in lipid metabolism [21], *bmm^1^* phenotypes were more pronounced (Figure S4A, S4B; Welch two-sample t-test, Kruskal-Wallis rank sum test). Defects in testis size were also observed at 14 days post-eclosion; suggesting testis size defects persist later into the life course (Figure S4C; Welch two-sample t-test). In contrast, the number of spermatid bundles per testis was not significantly different between *bmm^1^* and *bmm^rev^* males at this age (Figure S4D; Welch two-sample t-test), potentially due to a large decrease in the number of spermatid bundles in 14-day-old *bmm^rev^* males (Figure 4C, S4D). Together, these data suggest loss of *bmm* affects testis development and spermatogenesis. Similar phenotypes are observed in male mice without ATGL [52], and supplementing the diet of *bmm^1^* males with medium-chain triglycerides (MCT) partially rescues the testis and spermatogenic defects we observed in flies (Figure S4E, S4F; one-way ANOVA with Tukey multiple comparison test), as it does in mice [52,80]. This identifies similarities between flies and mice in fertility-related phenotypes associated with whole-body loss of *bmm*/*ATGL*.

**Figure 4.**
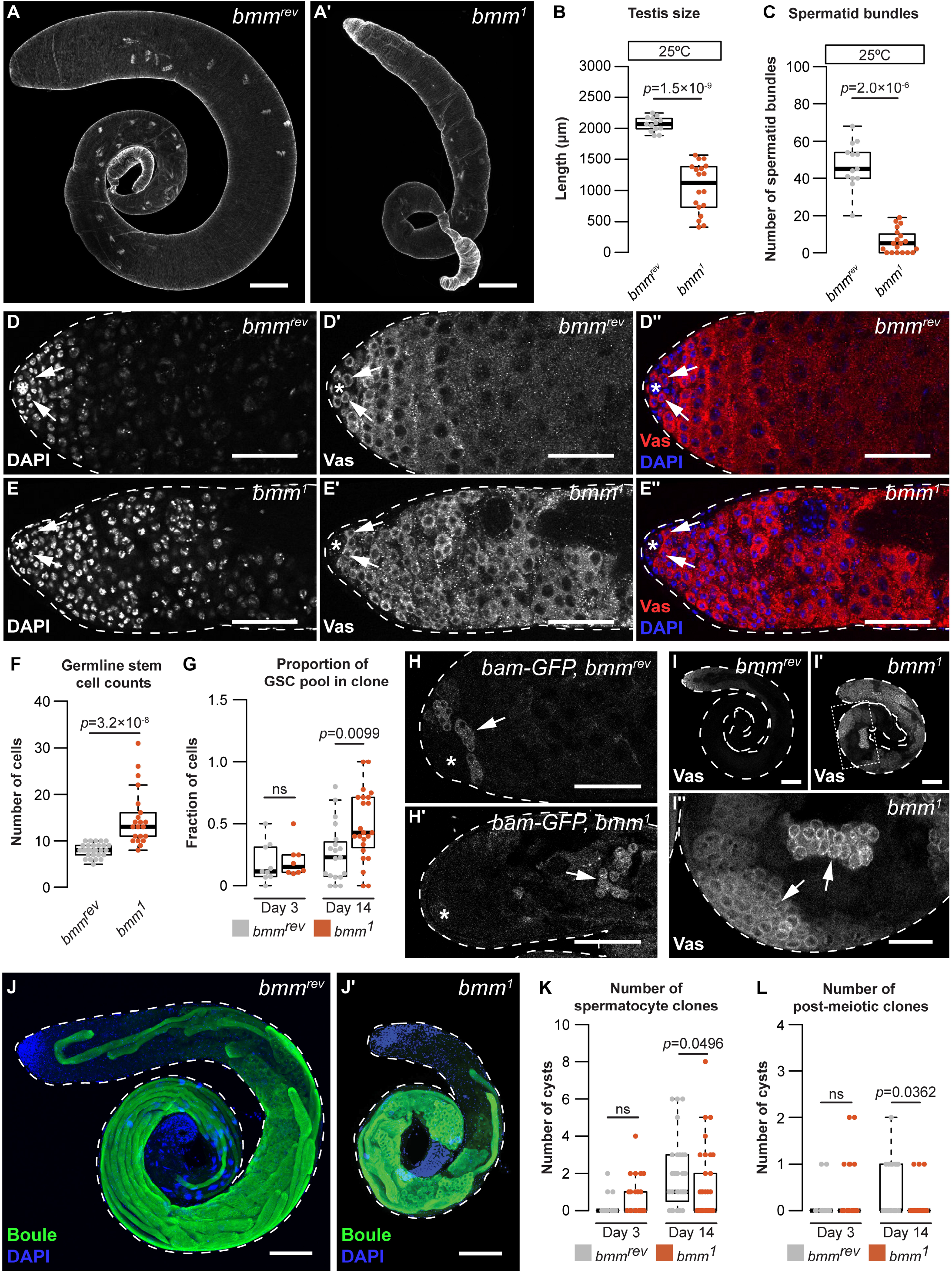
A cell-autonomous role for *bmm* in regulating spermatogenesis. Testes isolated from *bmm^rev^* (A) and *bmm^1^*(A’) animals raised at 25°C stained with phalloidin. Scale bars=100 μm. (B) Testis size in *bmm^1^* and *bmm^rev^* animals raised at 25°C. (C) Spermatid bundle number in *bmm^1^* and *bmm^rev^* testes from animals reared at 25°C. (D,E) Representative images of *bmm^rev^* (D) or *bmm^1^* (E) testes stained with DAPI and anti-Vasa antibody. Arrows indicate germline stem cells (GSC). Scale bar=50 μm. The hub is marked by an asterisk in all images. (F) GSC number in *bmm^1^* and *bmm^rev^* testes. (G) Proportion of GSCs that were either *bmm^1^* or *bmm^rev^*clones at 3 and 14 days post-clone induction. (H) Representative images of *bmm^rev^*(H) and *bmm^1^* (H’) testes carrying *bam-GFP*; data quantified in Figure S4J. Arrows indicate regions with high Bam-GFP. Scale bars=50 μm. (I) Representative images of *bmm^rev^* (I) or *bmm^1^*(I’,I’’) testes stained with anti-Vasa antibody. Arrows indicate Vasa-positive cysts in *bmm^1^* testis. Panel I’’ is magnified from the boxed region in I’. (I,I’) Scale bars=100 μm; (I’’) scale bar=50 μm. (J) Maximum projection of *bmm^rev^* (J) or *bmm^1^* (J’) testes stained with anti-Boule antibody (green) and DAPI (blue). Scale bars=100 μm. Number of *bmm^1^*and *bmm^rev^* spermatocyte clones (K) or post-meiotic clones (L) at 3 and 14 days post-clone induction. See also Supplemental Figure 4.

To explore spermatogenesis in *bmm^1^* animals, we used an antibody against the germline cell-specific marker Vasa to visualize the germline in the testes of *bmm^1^* and *bmm^rev^* males (Figure 4D, 4E) [81]. We observed a significant increase in the number of germline stem cells (GSC) (Figure 4F; Kruskal-Wallis rank sum test) and higher variability in GSC number in *bmm^1^* males (*p*=5.7×10^-12^ by F-test). Given that GSC number is affected by hub size and GSC proliferation [82,83], we monitored both parameters in *bmm^1^* and *bmm^rev^* controls. While hub size in *bmm^1^* testes was significantly larger than in testes from *bmm^rev^* controls (Figure S4G, S4H; Welch two-sample t-test), the number of phosphohistone H3-positive GSCs, which indicates proliferating GSCs, was unchanged in *bmm^1^* animals (Figure S4I; Kruskal-Wallis rank sum test). While this indicates a larger hub may partly explain *bmm’*s effect on GSC number, *bmm* also plays a cell-autonomous role in regulating GSCs, as we recovered a higher proportion of *bmm^1^* clones in the GSC pool compared with *bmm^rev^* clones at 14 days after clone induction (Figure 4G; Welch two-sample t-test). Given that we detected no effect of cell-autonomous *bmm* loss on the number of GSC LD (Figure 3C), more work will be needed to understand how *bmm* regulates GSCs at a stage prior to its effects on LD number. Future studies will also need to confirm whether *bmm^1^* mutant GSCs show an increased ability to occupy space at the hub.

Beyond GSCs, we uncovered additional spermatogenesis defects in *bmm^1^*testes. Peak Bam-GFP expression in germline cells of the testes from 0-day-old *bmm^1^* and *bmm^rev^* males showed GFP-positive cysts were significantly further away from the hub in *bmm^1^*testes (Figure 4H, S4J; Welch two-sample t-test). Indeed, 15/18 *bmm^1^* testes contained Vasa-positive cysts with large nuclei in the distal half of the testis (Figure 4I, arrows), a phenotype not present in *bmm^rev^* testes (0/8) (*p*=0.0005 by Pearson’s Chi-square test). Because these phenotypes are also seen in testes with differentiation defects [13,84], we recorded the stage of sperm development reached by the germline in *bmm^1^* testes. Most *bmm^1^* testes contained post-meiotic cells in males raised at 25°C (Figure S4K); however, germline development did not progress past the spermatocyte stage in most *bmm^1^* testes from animals raised at 29°C (Figure S4K). Testes from *bmm^1^* males reared at 25°C also had a smaller Boule-positive area (Figure 4J, S4L; Welch two-sample t-test), and fewer individualization complexes and waste bags (Figure S4M, S4N; Kruskal-Wallis rank sum test). Because Boule-positive area, individualization complexes, and waste bags are all markers for later stages in sperm development, these data indicate the loss of *bmm* causes a reduction in differentiated cell types. Because we observed significantly fewer *bmm^1^* spermatocyte and spermatid clones at 14 days after clone induction (Figure 4K, 4L; *p* = 0.0496, Kruskal-Wallis rank sum test), these effects on germline development may represent a cell-autonomous role for *bmm* in regulating spermatogenesis in this cell type. Given that the statistical significance of this finding was not as strong as for our other data, future studies should repeat this experiment with more samples. We also reveal a potential non-cell-autonomous role for somatic *bmm*. While there was no difference in the ratio of Zfh-1-positive cells between homozygous clones and heterozygous clones in animals carrying the *bmm^1^* or *bmm^rev^*alleles at 14 days post clone induction (Figure S4O; Kruskal-Wallis rank sum test), the distance from the hub to the Zfh-1 positive clones was significantly decreased in *bmm^1^* homozygous clones (Figure S4P; Kruskal-Wallis rank sum test). Together, these data indicate *bmm* may play a cell-autonomous role in germline cells, and potentially a non-cell-autonomous role in somatic cells, to regulate spermatogenesis.

### *brummer*-dependent regulation of testis triglyceride levels affects spermatogenesis

*ATGL* catalyzes the first and rate-limiting step of triglyceride hydrolysis [73,85,86]. Loss of this enzyme or its homologs leads to excess triglyceride accumulation [27,30,67,73,75] and shifts in multiple lipid classes [66,87–89]. To determine how loss of *bmm* affects spermatogenesis, we carried out whole-body mass spectrometry (MS)- based untargeted lipidomic profiling of *bmm^1^* and *bmm^rev^* males. Hierarchical clustering of lipid species suggests that *bmm^1^* and *bmm^rev^* males show distinct lipidomic profiles (Figure 5A). Overall, we detected 2464 and 1144 lipid features with high quantitative confidence in positive and negative ion modes, respectively. By matching experimental *m/z*, isotopic ratio, and tandem MS spectra to lipid libraries, we confirmed 293 unique lipid species (Supplemental table 1). We found 107 lipids had a significant change in abundance between *bmm^1^* and *bmm^rev^* males (p_adj_<0.05): 85 species were upregulated in *bmm^1^* males and 22 lipid species were downregulated. Among differentially regulated species from different lipid classes, triglyceride had the largest residual above expected proportion (*p*=5.00×10^-4^ by Pearson’s Chi-squared test). This suggests triglyceride is the lipid class most affected by loss of *bmm* (Figure 5B, 5C).

**Figure 5.**
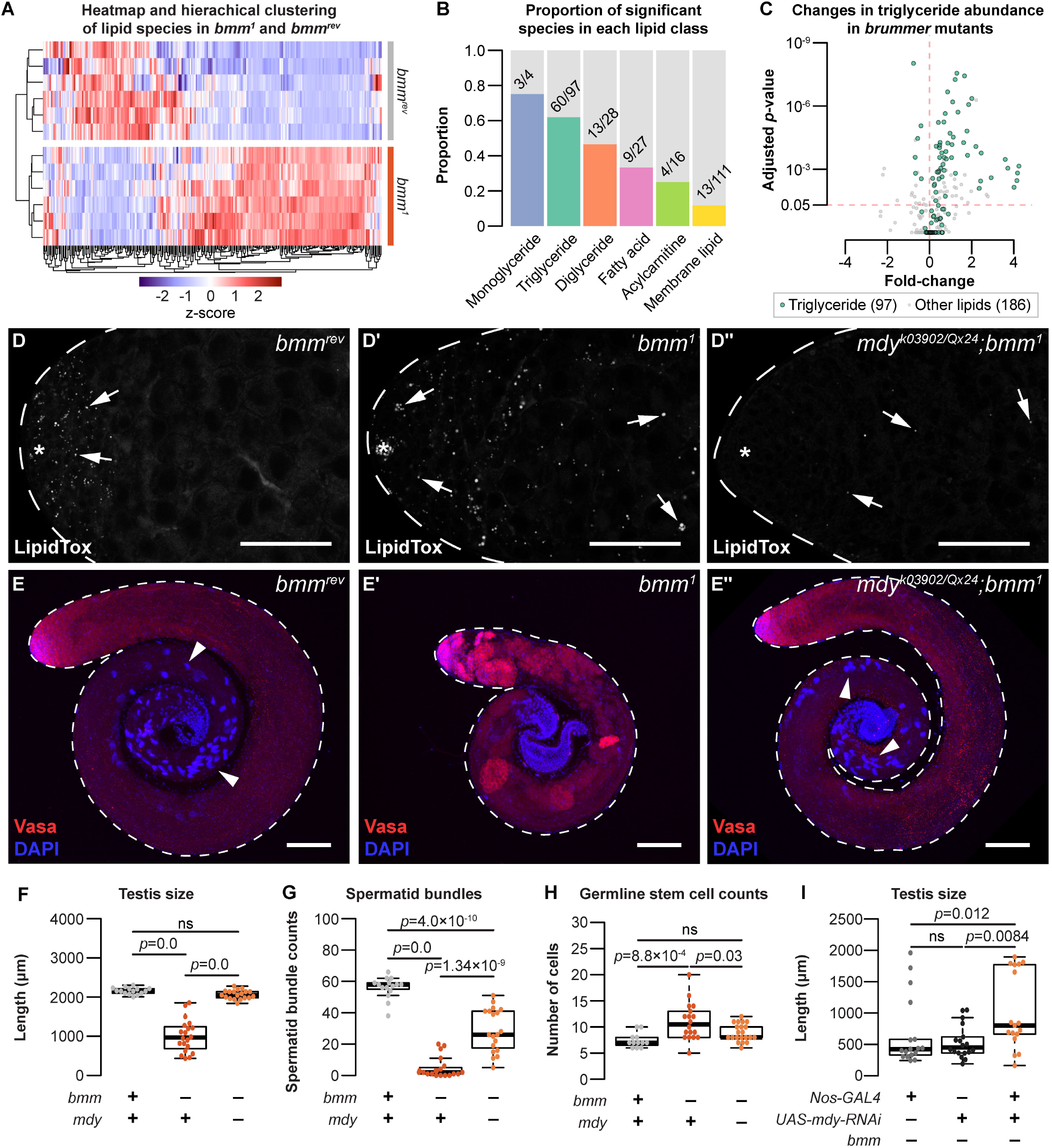
Loss of *bmm* disrupts triglyceride homeostasis and leads to spermatogenic defects. (A) Hierarchical clustering of lipid species detected in *bmm^rev^* and *bmm^1^* animals. (B) Histograms showing the proportion of significant species in each lipid class with different levels between *bmm^1^* and *bmm^rev^*. Numbers on histograms indicate the number of species with differences in abundance. (C) Volcano plot showing fold change in abundance of triglycerides (green; 97 species) and non-triglyceride lipids (grey; 186 species) in our dataset. (D) Arrows indicate testis LD stained with LipidTox Red in *bmm^rev^* (D), *bmm^1^* (D’), or *mdy^QX25/k03902^*; *bmm^1^* (D’’) animals. (E) Whole testes isolated from *bmm^rev^*(E), *bmm^1^* (E’), or *mdy^QX25/k03902^*;*bmm^1^*(E’’) animals stained with anti-Vasa antibody (red) and DAPI (blue). Arrowheads indicate spermatid bundles. Scale bars=100 μm. (F) Testis size in *bmm^rev^*, *bmm^1^*, and *mdy^QX25/k03902^*;*bmm^1^* animals. Spermatid bundles (G) and number of germline stem cells (H) in *bmm^rev^*, *bmm^1^*, and *mdy^QX25/k03902^*;*bmm^1^* animals. (I) Testis size in animals with germline-specific *mdy* knockdown (*nos-GAL4*>*mdy RNAi*; *bmm^1^*) compared with controls (*nos-GAL4*>*+*; *bmm^1^* and *+*>*mdy RNAi*; *bmm^1^*). See also Supplemental Figure 5.

In *bmm^1^* males, the majority of triglyceride species (55/97) were significantly higher in abundance compared with *bmm^rev^*control males. Because we observed a positive correlation between the fold increase in triglyceride abundance with both the number of double bonds (*p*=7.52×10^-8^ by Kendall’s rank correlation test; Figure S5A) and the number of carbons (*p*=2.77×10^-10^ by Kendall’s rank correlation test; Figure S5B), our data align well with *bmm*/*ATGL’s* known role in regulating triglyceride levels [67,68,73] and its substrate preference of long-chain polyunsaturated fatty acids [85]. While we also detected changes in species such as fatty acids, acylcarnitine, and membrane lipids (Figure S5C–S5H), in line with recent *Drosophila* lipidomic data [90,91], the striking accumulation of triglyceride in *bmm^1^* males suggested that excess testis triglyceride in *bmm^1^* males may contribute to their spermatogenic defects. To test this, we examined spermatogenesis in *bmm^1^* males carrying loss-of-function mutations in *midway* (*mdy*). *mdy* is the *Drosophila* homolog of *diacylglycerol O-acyltransferase 1* (*DGAT1*), and whole-body loss of *mdy* reduces whole-body triglyceride levels [92–94]. Importantly, testes isolated from males with global loss of both *bmm* and *mdy* (*mdy*^QX25/k03902^*;bmm^1^*) had fewer LD than testes dissected from *bmm^1^* males (Figures 5D, S5I; one-way ANOVA with Tukey multiple comparison test).

We found that testes isolated from *mdy^QX25/k03902^;bmm^1^*males were significantly larger and had more spermatid bundles than testes from *bmm^1^* males (Figure 5E–G; one-way ANOVA with Tukey multiple comparison test). The elevated number of GSCs in *bmm^1^* male testes was similarly rescued in *mdy^QX25/k03902^;bmm^1^* males (Figure 5H; one-way ANOVA with Tukey multiple comparison test). These data suggest that defective spermatogenesis in *bmm^1^* males can be partly attributed to excess triglyceride accumulation. Notably, at least some of the effects of global *mdy* loss on *bmm^1^* males can be attributed to the germline: RNAi-mediated knockdown of *mdy* in the germline of *bmm^1^* males partially rescued the defects in testis size (Figure 5I; Kruskal-Wallis rank sum test with Dunn’s multiple comparison test) and GSC variance (Figure S5J; *p*=4.5 x 10^-5^ and 8.2 x 10^-3^ by F-test from the GAL4- and UAS-only crosses, respectively). While future studies will need to test whether germline-specific loss of *mdy* also rescues spermatid number defects in *bmm^1^* males, our data suggest *bmm*-mediated regulation of testis triglyceride plays a previously unrecognized role in regulating sperm development.

## DISCUSSION

In this study, we used *Drosophila* to gain insight into how the neutral lipids, a major lipid class, contribute to sperm development. We describe the distribution of LD under normal physiological conditions in the *Drosophila* testis, and show that LD are present at the early stages of development in both somatic and germline cells. While many factors are known to regulate LD in nongonadal cell types, we reveal a cell-autonomous role for triglyceride lipase *bmm* in regulating testis LD during spermatogenesis. In particular, we identified a requirement for *bmm* in mediating the decrease in LD at the spermatogonia-spermatocyte transition. This regulation is important for sperm development, as our data indicates that loss of *bmm* causes a decrease in the number of differentiated cell types in the testis. This reduction in differentiated cell types may be attributed to a delay in differentiation, a block in differentiation, or to a loss of differentiated cells through cell death. Future studies will therefore be essential to resolve why *bmm* loss causes a reduction in differentiated cell types. Nevertheless, these defects in the number of differentiated cell types can be partially explained by the excess accumulation of triglyceride in flies lacking *bmm*, as global and cell type-specific inhibition of triglyceride synthesis rescues multiple spermatogenic defects in *bmm* mutants. Together, our data reveals previously unrecognized roles for LD and triglycerides during spermatogenesis, and for *bmm* as an important regulator of testis LD and germline development under normal physiological conditions.

One key outcome of our study was increased knowledge of LD regulation and function in the testis. Despite rapidly expanding knowledge of LD in cell types such as adipocytes or skeletal muscle, less is known about how LD influence spermatogenesis under normal physiological conditions. In mammals, testis LD contain cholesterol and play a role in promoting steroidogenesis [95,96]. In flies, we show that LD are present in the testis, and that excess accumulation of these LD affects sperm development. In nongonadal cell types, triglycerides provide a rich source of fatty acids for cellular ATP production, lipid building blocks to support membrane homeostasis and growth, and metabolites that can act as signaling molecules [26]. Because ATP production, lipid precursors, and lipid signaling all play roles in supporting normal sperm development [97,98], future studies will need to determine how each of these processes is affected when excess triglyceride accumulates in testis LD (Figure 6). It will also be important to determine whether it is the loss of metabolites produced by *bmm*’s enzymatic action, or an increase in triglycerides, that leads to the reduction in differentiated cell types during spermatogenesis. Together, these experiments will provide critical insight into how triglyceride stored within testis LD contributes to overall cellular lipid metabolism during spermatogenesis. Because of the parallel spermatogenic defects we observed in *bmm* mutants and *ATGL*-deficient mice, we expect these mechanisms will also operate in other species.

**Figure 6.**
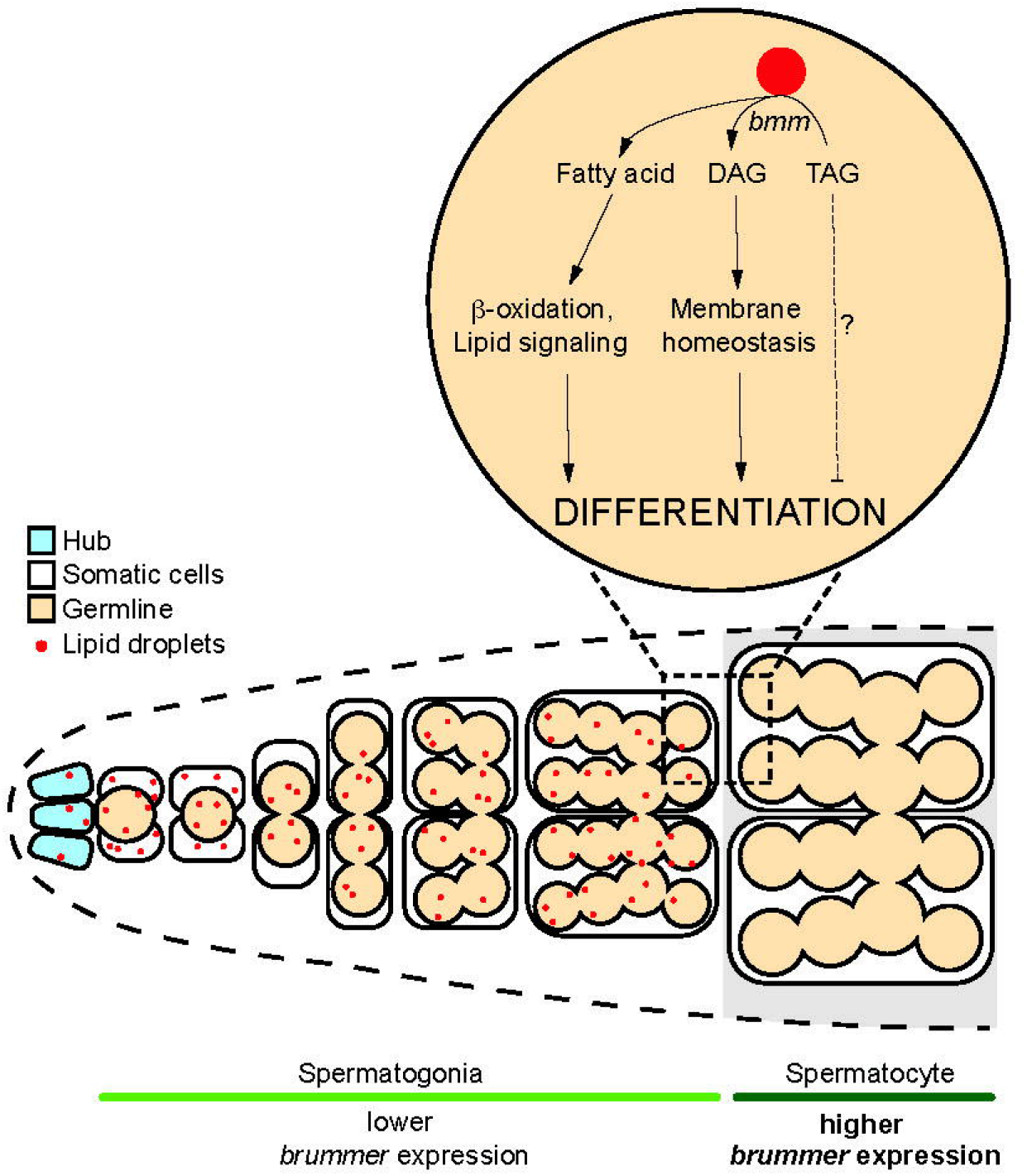
Model of *bmm*-mediated lipid droplet regulation in the *Drosophila* testis. Schematic representation summarizing *bmm*-mediated lipid droplet regulation in the testis during development.

A more comprehensive understanding of neutral lipid metabolism during sperm development will also emerge from studies on the upstream signaling networks that regulate testis LD and triglyceride. Given that we show an important and cell-autonomous role for *bmm* in regulating testis LD and triglyceride, future studies will need to identify factors that regulate *bmm* in the testis. Based on public single-cell RNAseq data and the *bmm-GFP* reporter strain, our data suggest *bmm* mRNA levels are differentially regulated between early and later stages of sperm development. Candidates for mediating this regulation include the insulin/insulin-like growth factor signaling pathway (IIS), Target of rapamycin (TOR) pathway, and nuclear factor κB/Relish pathway (NFκB), as all of these pathways influence *bmm* mRNA levels in nongonadal cell types [99–105]. Beyond mRNA levels, Bmm protein levels and post-translational modifications may also be differentially regulating during spermatogenesis. For example, studies show that the proteins encoded by *bmm* homologs in other animals are regulated by phosphorylation [106], mediated by kinases such as adenosine monophosphate-activated protein kinase (AMPK) and protein kinase A (PKA) [107–109]. Importantly, many of these pathways, including IIS, TOR, AMPK, NFκB and possibly PKA influence *Drosophila* sperm development [110–115]. Identifying the signaling networks that influence *bmm* regulation during sperm development will therefore lead to a deeper understanding of how testis LD and triglyceride are coordinated with physiological factors to promote normal spermatogenesis. Because pathways such as IIS and AMPK, and others, regulate sperm development in other species [116–118], these insights may reveal conserved mechanisms that govern the regulation of cellular neutral lipid metabolism during sperm development.

## ACKNOWLEDGEMENTS

We thank Dr. Ronald Kühnlein for *bmm^1^*and *bmm^rev^* lines [67], Dr. Michael Welte for *UAS-GFP-LD* [58], and Dr. Kaeko Kamei for *bmm-GFP* [77]. We used stocks from the Bloomington *Drosophila* Stock Center (NIH P40OD018537) and Vienna *Drosophila* Resource Center (VDRC). We acknowledge critical resources and information provided by FlyBase [119] (supported by the National Human Genome Research Institute at the U.S. National Institutes of Health (<u>U41 HG000739</u>) and the British Medical Research Council (<u>MR/N030117/1</u>)). This work was supported by the Life Sciences Institutes Imaging Core, supported by the UBC GREx Biological Resilience Initiative. Funding for this study was provided by grants to EJR from the Canadian Institutes for Health Research (PJT-153072), Michael Smith Foundation for Health Research (16876), and the Canadian Foundation for Innovation (JELF-34879). GT was supported by a grant from the Natural Sciences and Engineering Research Council (NSERC; 2018-04648), TH/HY/CW were supported by NSERC (2020-04895), MA was supported by the Jacob’s foundation. We would like to acknowledge that our research takes place on the traditional, ancestral, and unceded territory of the Musqueam people; a privilege for which we are grateful.

## AUTHOR CONTRIBUTIONS

Conceptualization, C.C. and E.J.R.; Methodology, C.C. and Y.Y.P.; Software, C.C.; Investigation, C.C., H.Y., and Y.Y.P.; Lipidomics, M.A., H.Y., C.W., T.H.; Writing – Original Draft, C.C. and E.J.R.; Writing – Review and Editing, C.C., E.J.R., and Y.Y.P.; Supervision, E.J.R., G.T., and T.H.; Project administration, E.J.R.; Funding Acquisition, E.J.R., G.T., and T.H.

## MATERIALS AND METHODS

### Materials and Resource availability

*Drosophila* strains and their source are listed in a Key Resources table. Further information and requests for resources and reagents should be directed to, and will be fulfilled by, lead contact Dr. Elizabeth J. Rideout.

### Data and Code availability

All raw data and results of statistical tests reported in this paper are located in Supplementary files 1-4. This paper does not report original code. Any additional information required to reanalyze the data reported in this paper is available from the lead contact upon request.

### Fly husbandry

Fly stocks were maintained at room temperature in 12:12 hour light:dark cycle. Unless otherwise indicated, all flies were raised at 25°C with a density of 50 larvae per 10 mL fly media. Because this project examines sperm development, we used male flies in all experiments. Fly media contained 20.5 g sucrose (SU10, Snow Cap), 70.9 g Dextrose (SUG8, Snow Cap), 48.5 g cornmeal (AO18006, Snow Cap), 30.3 g baker’s yeast (NB10, Snow Cap), 4.55 g agar (DR-820-25F, SciMart), 0.5 g calcium chloride dihydrate (CCL302.1, BioShop Canada), 0.5 g magnesium sulfate heptahydrate (MAG511.1, BioShop Canada), 4.9 mL propionic acids (P1386, Sigma-Aldrich), and 488 μL phosphoric acid (P5811, Sigma-Aldrich) per 1L of media. For diets with medium- or long-chain triglyceride, 4 g of coconut oil (medium chain triglyceride) or olive oil (long chain triglyceride) was added per 100 mL of media described above prior to cooling. Males were collected and dissected within 24 hours of eclosion unless otherwise indicated. Fixations were performed at room temperature with 4% paraformaldehyde (CA11021-168, VWR) in PBS for 20 minutes on a rotating platform followed by washing in PBS twice before staining. Fly strains used in our study are listed in a Key Resources table.

### Testis cell stage classification and measurements

Cells at an early stage of development (stem cells and early-stage somatic and germline cells) were located in the apical region of the testis, and were identified by their small and dense nuclei[120]. GSCs were defined as Vasa-positive cells in direct contact with the hub; proliferating GSCs were identified as Vasa-positive cells in direct contact with the hub that were also phospho-H3 positive. Cells in the testis region occupied by primary spermatocytes were identified by their large cell size and decondensed chromosome staining occupying three nuclear domains [120]. Spermatid bundles were identified by their condensed and needle-shaped nuclei, which roughly corresponds to nuclei with protamine-based chromatin [121]. The hub was identified as the FasIII-positive area of the testis. Hub size was estimated by measuring the FasIII-positive area in a Z-projected image of the hub in each testis. Z-projections were made using the ‘sum slices’ function in Fiji. Testis size was measured by quantifying the length of a line drawn down the middle of a testis image; starting from the apical tip of the testis and ending where the testis meets the seminal vesicle.

### FLP-FRT clone induction

Adult males were collected at 3-5 days post-eclosion and heat-shocked three times at 37°C for 30 min followed by a 10 min rest period at room temperature between heat shocks. After heat-shock, the flies were incubated at room temperature until dissection.

### Immunohistochemistry

Fixed samples were rinsed three times with blocking solution containing 0.2% bovine serum albumin (A4503, Sigma-Aldrich), 0.3% Triton-X in PBS, then blocked for 1 hr on a rotating platform at room temperature. During the incubation, the blocking solution was changed every 15 minutes. After blocking, the sample were resuspended in blocking solution with the appropriate concentration of primary antibody (see Key Resources table), and incubated overnight at 4°C. Samples were rinsed three times with blocking solution after removing primary antibody, and blocked for one hour on a rotating platform in blocking solution. Secondary antibody was applied in blocking solution and left on the rotating platform at room temperature for 40 min. The sample was rinsed with blocking solution three more times, and washed four times for 15 min per wash in blocking solution. Testis samples were resuspended in Vectashield mounting media with DAPI (H-1200-10, Vector Laboratory) or SlowFade Diamond mounting media (S36972, Thermo Fisher Scientific) prior to mounting.

### Lipid droplet staining

Fixed testes were briefly permeabilized with 0.1% Triton-X in PBS for 5 min prior to applying phalloidin. For BODIPY (4,4-Difluoro-1,3,5,7,8- Pentamethyl-4-Bora-3a,4a-Diaza-*s*-Indacene) staining, samples were suspended in PBS containing 10 μg/mL DAPI (2879083-5mg, PeproTech), 1:500 BODIPY 495/503 (Thermo Fisher Scientific D3922), and 1:1000 phalloidin iFluor647 (ab176759, Abcam) or 1:40 phalloidin TexasRed (T7471, Thermo Fisher Scientific). For staining with LipidTox Red, samples were suspended in PBS containing 10 μg/mL DAPI (2879083-5mg, PeproTech), 1:200 LipidTox Red (H34476, Thermo Fisher Scientific), and 1:1000 phalloidin iFluor647 (ab176759, Abcam). For staining free sterols, samples were prepared as for BODIPY staining with 50 μg/mL filipin in place of BODIPY for 30 min. Samples were incubated on a rotating platform for 40 minutes at room temperature. After incubation, samples were washed twice with PBS, then resuspended in SlowFade Diamond mounting media (Thermo Fisher Scientific S36972) prior to mounting.

### Image acquisition and processing

All images were acquired on a Leica SP5 confocal microscope system with 20X or 40X objectives and quantified with Fiji image analysis software[122].

### Drosophila lipidomics

*Drosophila* extracts were prepared following the previously reported protocol[123]. Briefly, 10 *Drosophila* males (∼10 mg) were weighed, 300 µL of ice-cold methanol/water mixture (9:1, v:v) was added to these males, and the samples were homogenized with glass beads using a bead beater (mini-beadbeater-16, BioSpec, Bartlesville, Ok, USA). Sample weight was used for sample normalization. Fly lysate was kept at -20°C for 4 hours for protein precipitation. Then, 900 µL of methyl tert-butyl ether was added and the solution was shaken for 5 min to extract lipids. To induce phase separation 285 µL of water was added, followed by centrifugation. The upper layer was separated, dried, and reconstituted in isopropanol/acetonitrile (1:1, v:v) for liquid chromatography-mass spectrometry (LC-MS) analysis. The volume of reconstitution solution was proportional to sample weight for normalization. Quality control (QC) samples were prepared by pooling 20 μL aliquot from each sample. The method blank sample was prepared using an identical workflow but without adding *Drosophila*.

*Drosophila* extracts were analyzed on an UHR-QqTOF (Ultra-High Resolution Qq-Time-Of-Flight) mass spectrometry Impact II (Bruker Daltonics, Bremen, Germany) interfaced with an Agilent 1290 Infinity II LC Systems (Agilent Technologies, Santa Clara, CA, USA). LC separation was performed using a Waters reversed-phase (RP) UPLC Acquity BEH C18 Column (1.7 µm, 1.0 mm ×100 mm, 130 Å) (Milford, MA, USA) maintained at 30°C. For positive ion mode, the mobile phase A was 60% acetonitrile in water and the mobile phase B was 90% isopropanol in acetonitrile, both containing 5 mM ammonium formate (pH = 4.8, adjusted by formic acid). For negative ion mode, the mobile phase A was 60% acetonitrile in water and the mobile phase B was 90% isopropanol in acetonitrile, both containing 5 mM NH_4_FA (pH = 9.8, adjusted by ammonium hydroxide). The LC gradient for positive and negative ion modes was set as follows: 0 min, 5% B; 8 min, 40% B; 14 min, 70% B; 20 min, 95% B; 23 min, 95% B; 24 min, 5% B; 33 min, 5% B. The flow rate was 0.1 mL/min. The injection volume was optimized to 2 µL in positive mode and 5 µL in negative mode using QC sample. The ESI source conditions were set as follows: dry gas temperature, 220 °C; dry gas flow, 7 L/min; nebulizer gas pressure, 1.6 bar; capillary voltage, 4500 V for positive mode and 3000 V for negative mode. The MS1 analysis was conducted using following parameters: mass range, 70-1000 m/z; spectrum type: centroid, calculated using maximum intensity; absolute intensity threshold: 250. Data-dependent MS/MS analysis parameters: collision energy: 16-30 eV; cycle time, 3 s; spectra rate: 4 Hz when intensity < 10^4^ and 12 Hz when intensity > 10^5^, linearly increased from 10^4^ to 10^5^. External calibration was applied using sodium formate to ensure the m/z accuracy before sample analysis.

The raw LC-MS data were processed using MS-DIAL (ver. 4.38)[124]. The detailed MS-DIAL parameters are: MS1 tolerance, 0.01 Da; MS/MS tolerance, 0.05; mass slice width, 0.05 Da; smoothing method, linear weighted moving average; smoothing level, 3 scans; minimum peak width, 5 scans. Lipid features with high quantitative confidence were selected by the following criteria: retention time was within the gradient elution time (< 23 min); average intensity in QC samples is larger than 5-fold of the intensity in method blank sample. Lipid identification was performed by matching experimental precursor m/z, isotopic ratio and MS/MS spectrum against the LipidBlast libraries embedded in MS-DIAL. To improve the quantification accuracy, the measured MS signal intensities were corrected using serial diluted QC samples following the reported workflow[125].

#### Quantification and statistical analysis

All microscopy images were quantified using Fiji software[122]. For lipid droplet counts, a single optical slice through the middle of the testis containing the hub was used with the exception of FLP-FRT experiment where all lipid droplets within a GFP-negative cyst were counted (Figure 2I). All statistical analyses were done using R (obtained from https://cran.r-project.org). With exception of data concerning spatial distribution, and lipidomic data, Shapiro-Wilk test (via *shapiro.test* in base R) was used to assess normality of distribution prior to testing for significance. Kruskal-Wallis rank sum test (from the R package *coin*) and Dunn’s test (from the R package *dunn.test*) were used in place of Welch two-sample t-test and Tukey’s multiple comparison test when the assumption of normality was not met. For testing differences in variance between two populations, F-test (via *var.test* in base R) was used. For testing differences in spatial distribution, two-sample Kolmogorov-Smirnov test (via *ks.test* in base R) was used. All *p*-values are indicated in figures; extremely small *p*-values are listed as *p*<2.2 x 10^-16^.

## RESOURCE TABLE

| REAGENT or RESOURCE | SOURCE | RESOURCE # |
| --- | --- | --- |
| **Antibodies** |  |  |
| Anti-Vasa (1:200) | Gift from Dr. R. Lehman,<br>MIT |  |
| Anti-Eya (1:50) | Developmental Studies<br>Hybridoma Bank<br>(DSHB) | eya10H6 |
| Anti-zfh1 (1:1000) | Gift from Dr. J. Skeath,<br>WUSTL |  |
| Anti-boule (1:1000) | Gift from Dr. S.<br>Wasserman, UCSD |  |
| Anti-phospho-histone H3 (1:1000) | Millipore Sigma | 05-1354 |
| **Experimental models: *Drosophila melanogaster*** |  |  |
| *w*<sup>1118</sup> | Bloomington *Drosophila*<br>stock center | 3605 |
| *CantonS* | Bloomington *Drosophila*<br>stock center | 64349 |
| *OregonR* | Bloomington *Drosophila*<br>stock center | 25211 |
| *bmm*<sup>1</sup> | Gift from Dr. R. Kühnlein |  |
| *bmm*<sup>rev</sup> | Gift from Dr. R. Kühnlein |  |
| *mdy*[Qx25], *cn*[1], *bw*[1]/CyO, *l*(2)*DTS513*[1] | Bloomington *Drosophila*<br>stock center | 5095 |
| *y*[1], *w*[67c23]; *P*{*lacW*}*Cse*1[*k03802*], *mdy*[*k03902*]/CyO | Bloomington *Drosophila*<br>stock center | 10536 |
| *w*[1118]; *P*{*GD1749*}v6367 (*UAS-mdy-RNAi*) | Vienna *Drosophila*<br>resource center | 6367 |
| *nos-GAL4::VP16* | Bloomington *Drosophila*<br>stock center | 7303 |
| *Tj-GAL4* | Gift from Dr. D. Godt,<br>University of Toronto |  |
| *c587-GAL4* | Bloomington *Drosophila*<br>stock center | 67747 |
| Bam-GFP | <sup>40</sup> |  |
| *bmm-GFP* | Gift from Dr. K. Kamei<sup>57</sup> |  |
| *GFP-LD* | Gift from Dr. M. Welte<sup>36</sup> |  |
| *P{neoFRT}82B, bmm[1]* | This study |  |
| *P{neoFRT}82B, bmm[rev]* | This study |  |
| *bam-GFP, bmm[1]* | This study |  |
| *bam-GFP, bmm[rev]* | This study |  |
| **Software and algorithms** |  |  |
| Fiji | https://imagej.net/software/fiji/ |  |
| R | https://cran.r-project.org |  |

## SUPPLEMENTAL TABLE AND FILES

**Supplemental table 1** – Table showing identified lipid species from untargeted lipidomic analysis.

**Supplementary file 1** – Raw data and statistical outputs from Figure 1.

**Supplementary file 2** – Raw data and statistical outputs from Figure 2 and Figure 3.

**Supplementary file 3** – Raw data and statistical outputs from Figure 4 and Supplemental figure 4.

**Supplementary file 4** – Raw data and statistical outputs from Figure 5 and Supplemental figure 5.

## SUPPLEMENTAL FIGURE LEGENDS

**Supplemental Figure 1 related to Figure 1.**
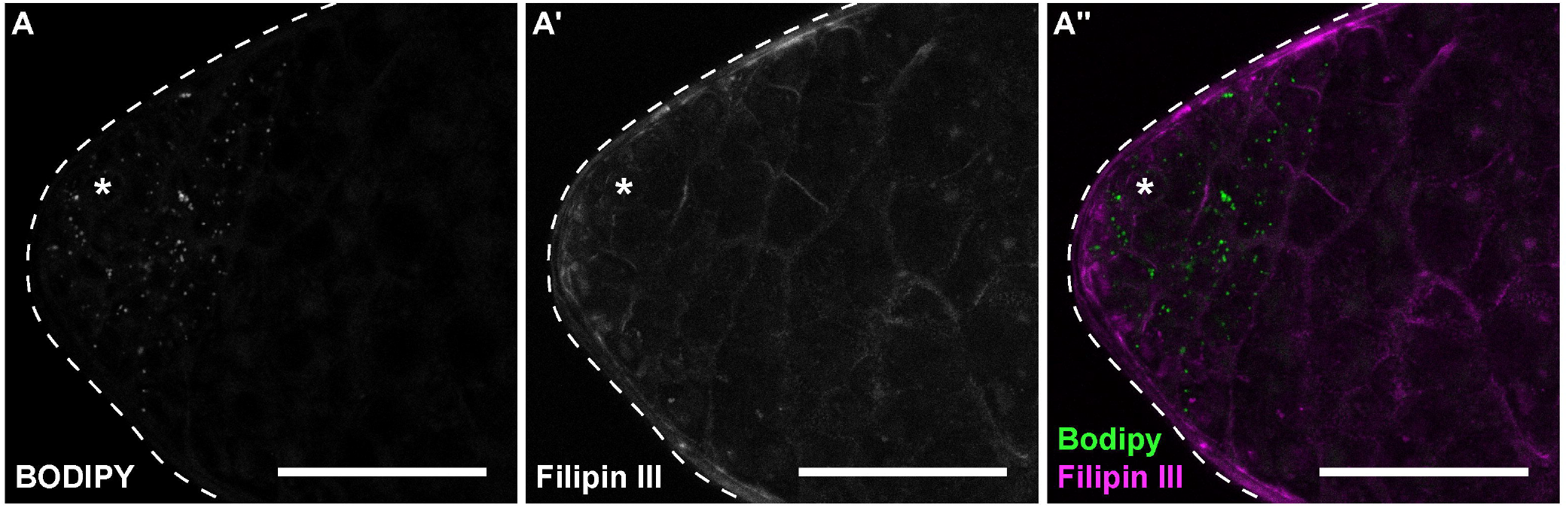
Cholesterol is absent from testis lipid droplets. (A) Testes stained with BODIPY (A) to detect neutral lipids and Filipin III (A’) to detect free cholesterol. Scale bars=50 μm.

**Supplemental Figure 2 related to Figure 2.**
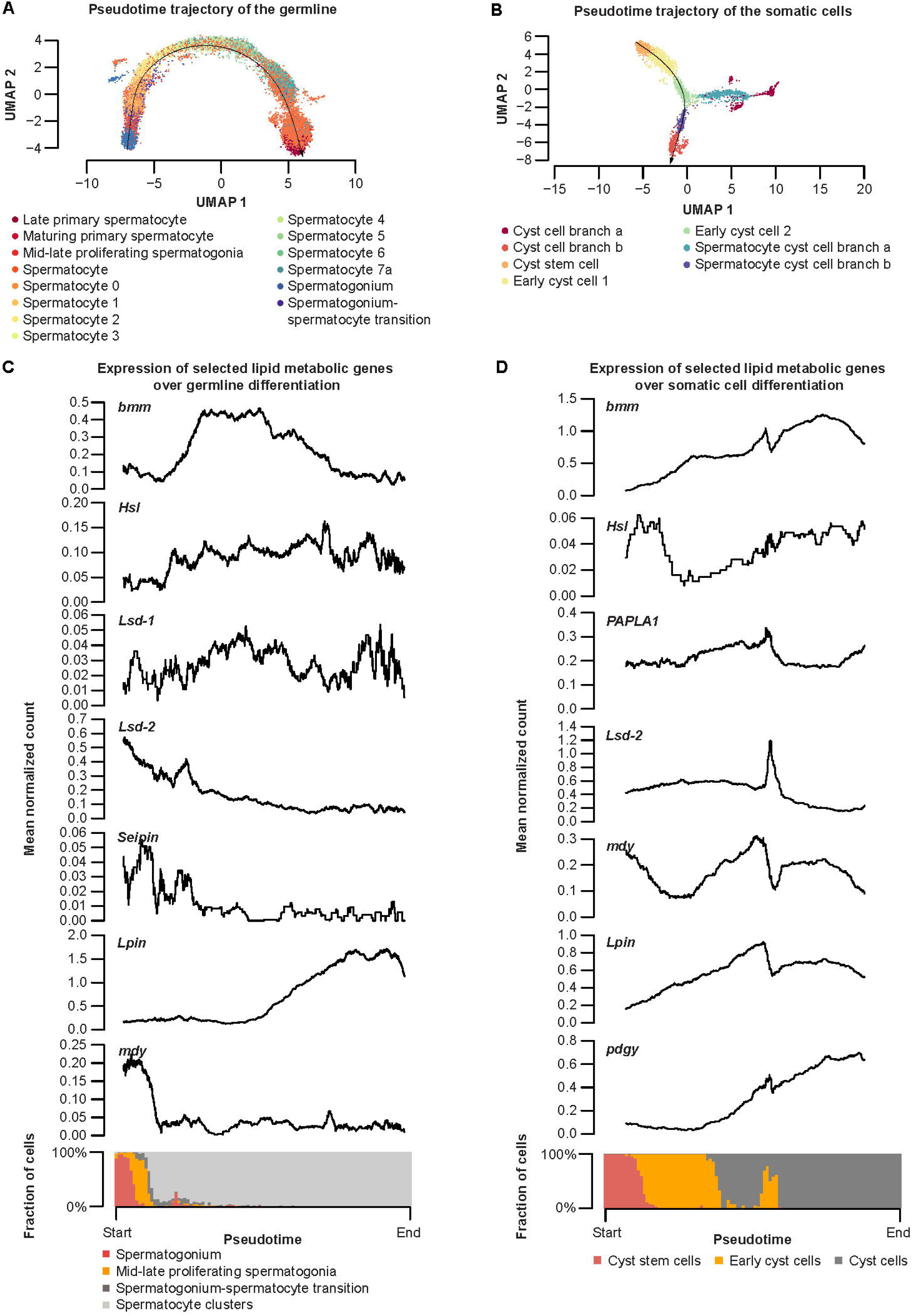
Expression of *brummer* and selected lipid metabolic genes during spermatogenesis in germline and somatic lineages. (A) Pseudotime trajectory of germline (black line) based on single-cell RNA sequencing data [78]. Individual cells are labeled according to the annotation within the data set. (B) Pseudotime trajectory of the somatic cells (black line) based on publicly available single-cell RNA sequencing data [78]. Individual cells are labeled according to the annotation within the data set. Only one trajectory (branch b) is marked and used for panel D. (C) Rolling average of normalized transcript counts in the germline along the trajectory shown in panel A are plotted as a black line on the upper panel. Composition of cell types mapped on to the trajectory at each time point is shown at the bottom of panel B. (D) Rolling average of normalized transcript counts in somatic cells plotted as a black line along the trajectory shown in C (upper panel). Composition of cell types mapped on to the trajectory at each time point is shown at the bottom of panel D.

**Supplemental Figure 3 related to Figure 2.**
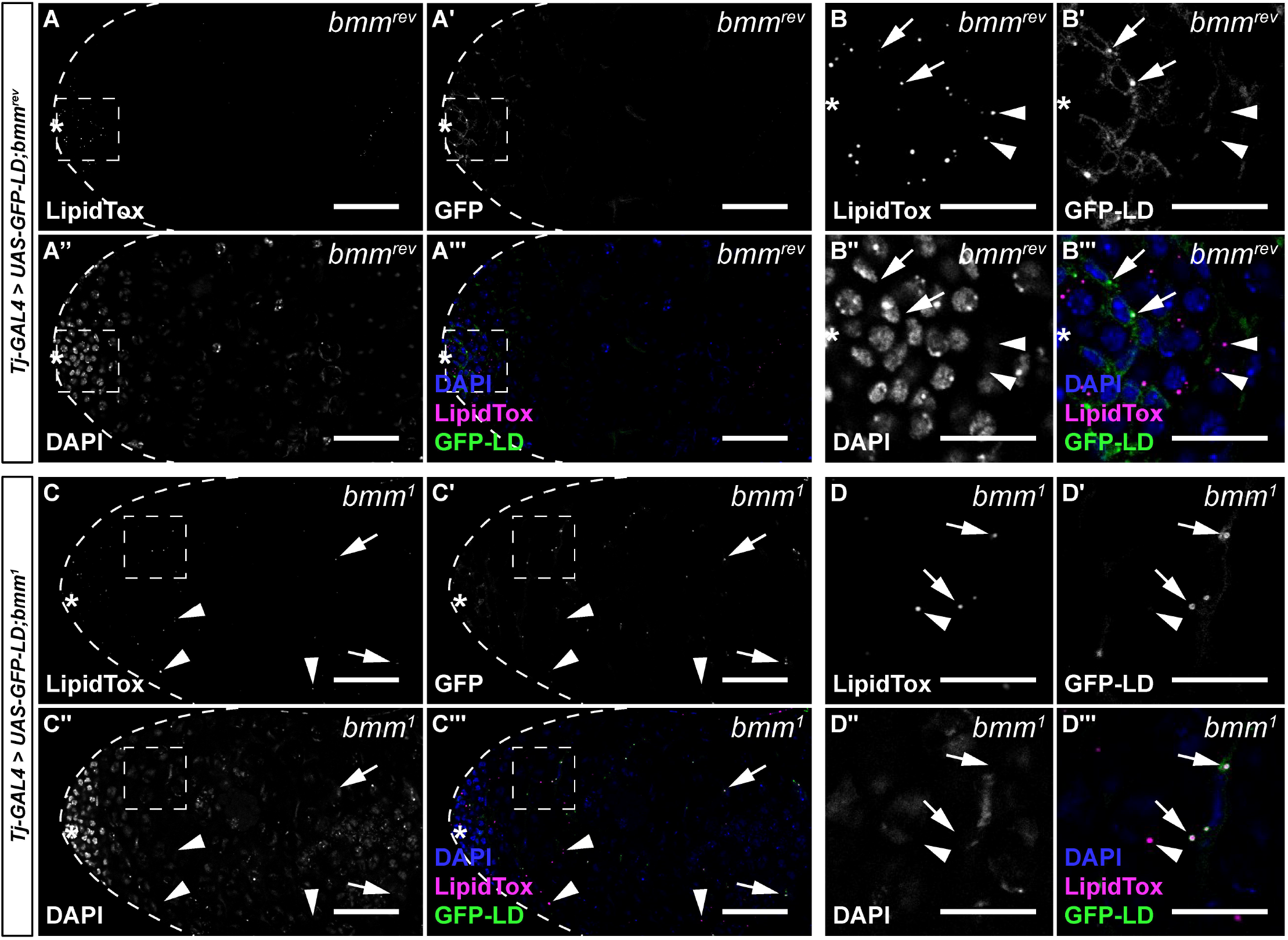
*brummer* regulates lipid droplets in both germline and somatic cells of the testis. (A-D) Representative images of *bmm^rev^* (A and B) and *bmm^1^*(C and D) testes with somatic over-expression of *GFP-LD* (*Tj-GAL4*>*UAS-GFP-LD*). Panel F and H contain magnified images of the area indicated by the boxes in panel A and C, respectively. In *bmm^rev^* testes, LD were restricted to a region near the apical tip (A) of the testis in both somatic (B–B’’’ arrows) and germline cells (B–B’’’ arrowheads). In *bmm^1^*testes, LD were present in both somatic (C–D arrows) and germline cells (C–D arrowheads), near the apical tip of the testis in a region corresponding to early-stage germ cells and in the region corresponding to spermatocytes. (A,C) Scale bars=50 μm; (B,D) scale bars=20 μm.

**Supplemental Figure 4 related to Figure 4.**
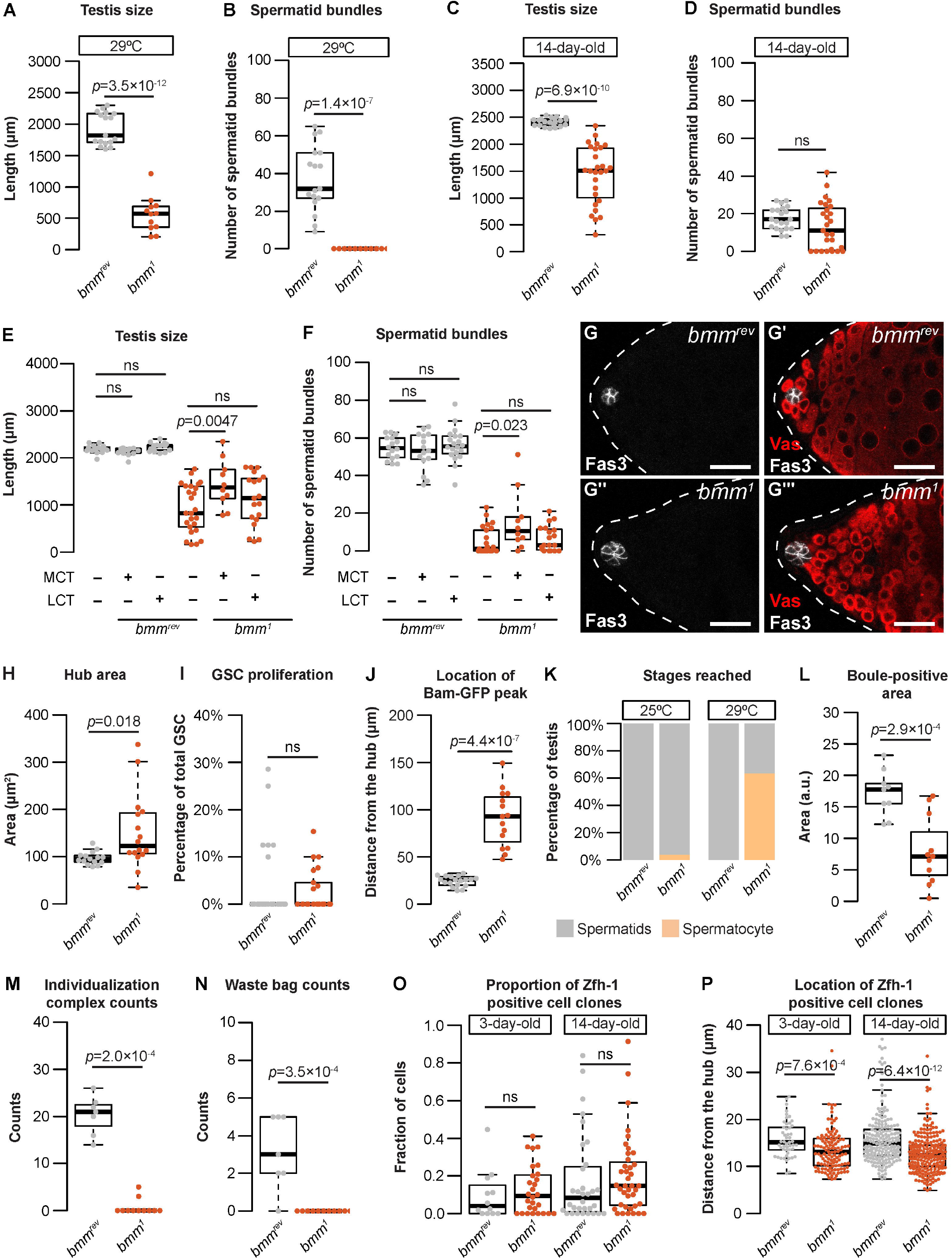
Additional characterization of testis development and spermatogenesis in animals lacking *bmm*. (A) Testis size was smaller in *bmm^1^* mutant animals compared with *bmm^rev^* controls at <24 hr post-eclosion when raised at 29°C (A; Welch two-sample t-test). (B) The number of spermatid bundles was significantly lower in *bmm^1^*mutant animals compared with *bmm^rev^* controls at <24 hr post-eclosion when raised at 29°C (Kruskal-Wallis rank sum test). (C) Testis size was significantly smaller in *bmm^1^* mutant males compared with *bmm^rev^*control males at 14-days post-eclosion (Welch two-sample t-test). (D) While the median number of spermatid bundles was not significantly different between *bmm^1^* mutant males and *bmm^rev^* control males at 14 days post-eclosion (Welch two-sample t-test), 8/27 *bmm^1^*testis had no spermatid bundles, a phenotype absent in age-matched *bmm^rev^*males (0/22) (p=0.0163, Pearson’s Chi-squared test), suggesting a subtle defect is present. (E) Food supplemented with 4% medium chain triglyceride (MCT), but not long chain triglyceride (LCT), significantly increased testis length in *bmm^1^* animals but had no effect on this phenotype in *bmm^rev^* control animals (one-way ANOVA with Tukey multiple comparison test). (F) Food supplemented with 4% medium chain triglyceride significantly increased the number of spermatid bundles in *bmm^1^*testes but had no effect on this phenotype in *bmm^rev^* control animals (one-way ANOVA with Tukey multiple comparison test). (G) Representative images of *bmm^rev^* (G–G’) or *bmm^1^* (G’’– G’’’) testes stained for Fas3 (G and G’’) and Vas (G’ and G’’’). Scale bars=25 μm. (H) Quantification of hub area in *bmm^rev^*or *bmm^1^* testes showed a significantly larger hub size in *bmm^1^*testes (Welch two-sample t-test). (I) The number of germline stem cells (GSC) undergoing mitosis (phospho-histone H3^+^ GSC/total GSC) was not significantly different between *bmm^1^* and *bmm^rev^*testes (Kruskal-Wallis rank sum test). (J) The distance between the hub and the first Bam-GFP positive cyst (Figure 3H) was significantly higher in *bmm^1^* testes than in *bmm^rev^*testes (Welch two-sample t-test). (K) All *bmm^rev^* testes and most *bmm^1^* testes contained spermatids when raised at 25°C; however, the most advanced stage of spermatogenesis observed in the majority of *bmm^1^* testes isolated from animals reared at 29°C was the spermatocyte stage. (L) Testes isolated from *bmm^1^* animals showed a significantly smaller Boule-positive area than control testes (Welch two-sample t-test). (M) Testes isolated from *bmm^1^* animals contain fewer individualization complexes than *bmm^rev^* control testes (Kruskal-Wallis rank sum test). (N) Fewer waste bags were present in testes isolated from *bmm^1^*animals compared with *bmm^rev^* control testes (Kruskal-Wallis rank sum test). (O) Proportion of Zfh-1 positive clones that were homozygous for either *bmm^1^* or *bmm^rev^* at 3 and 14 days post-clone induction (Kruskal-Wallis rank sum test). (P) The location of Zfh-1 positive clones homozygous for either *bmm^1^* or *bmm^rev^* at 3 and 14 days post-clone induction measured as the distance from the center of the hub. Clones homozygous for *bmm^1^* locate significantly closer to the hub than the wildtype clones (Kruskal-Wallis rank sum test).

**Supplemental Figure 5 related to Figure 5.**
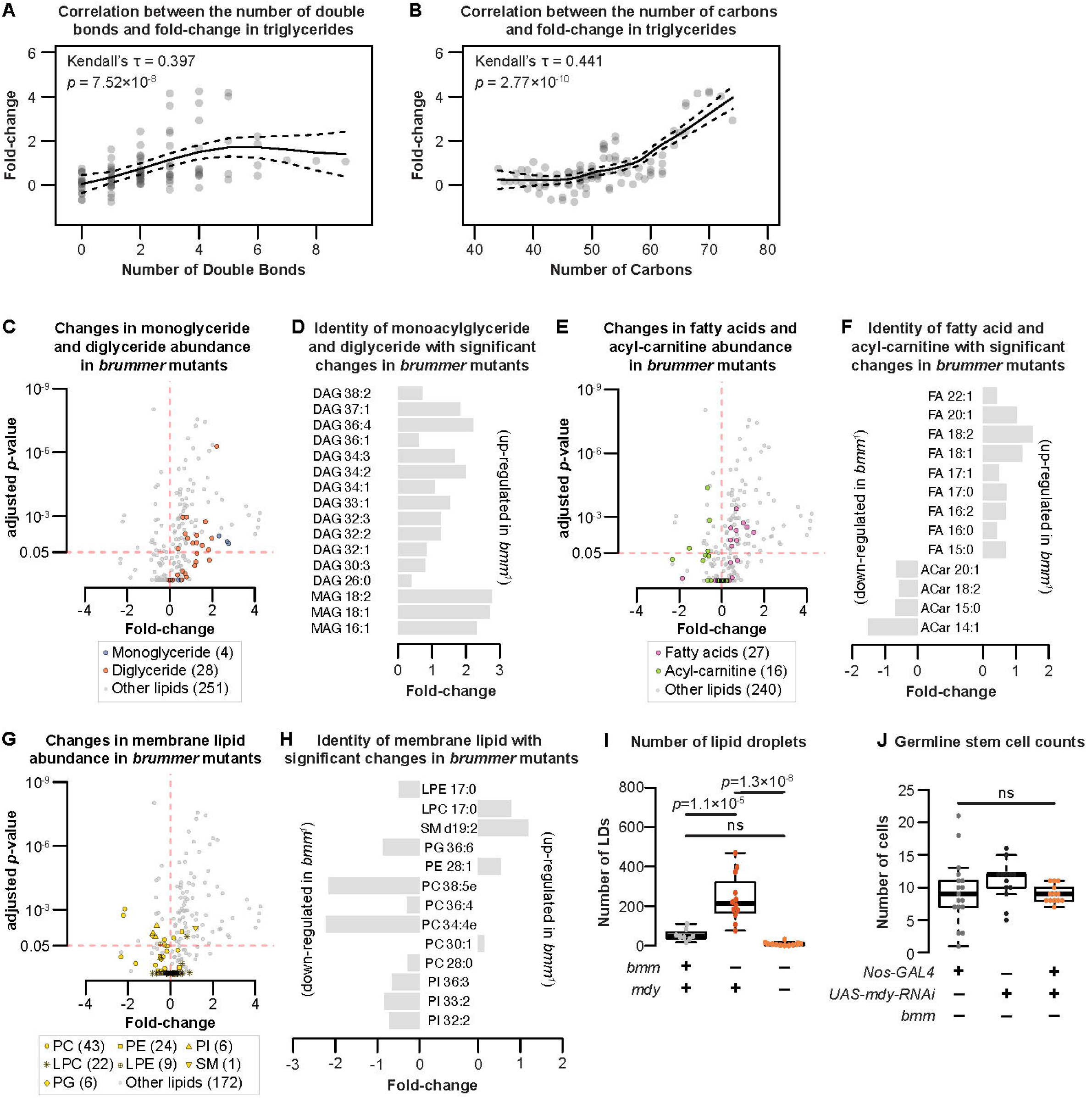
Lipidomic analysis of animals lacking *bmm*. (A) Higher fold-changes of triglycerides in *bmm^1^* animals were associated with less saturation in the acyl-groups (Kendall’s rank correlation test). (B) Higher fold-changes of triglycerides in *bmm^1^* animals were associated with higher number of carbons in the acyl-groups (Kendall’s rank correlation test). Each dot represents a single triglyceride species for panel B and C. (C) Volcano plot of identified lipids; monoglycerides shown in blue and diglycerides shown in orange. Many monoglycerides and diglycerides show increase in fold-change in *bmm^1^*males. (D) The number of carbon and the degree of saturation of monoglycerides (MAG) and diglycerides (DAG) with significant changes in abundance between *bmm^1^* and *bmm^rev^* males. (E) Volcano plot of identified lipids; fatty acids shown in magenta and acyl-carnitine shown in green. Many fatty acids show an increase in fold-change while many acyl-carnitines show a decrease in fold-change in *bmm^1^*males. (F) The number of carbon and the degree of saturation of fatty acids (FA) and acyl-carnitines (ACar) with significant changes in abundance between *bmm^1^*and *bmm^rev^* males. (G) Volcano plot of identified lipids; membrane lipids shown in yellow. (H) The number of carbon and the degree of saturation of membrane lipids with significant changes in abundance between *bmm^1^* and *bmm^rev^* males. For panel G and H, PC: phosphatidylcholine; PE: phosphatidylethanolamine; PI: phosphatidylinositol; LPC: lysophosphatidylcholine; LPE: lysophosphatidylethanolamine; SM: sphingomyelin; PG: phosphatidylglycerol. (I) Loss of *mdy* function rescued the elevated number of LD in *bmm^1^* testes to control levels (one-way ANOVA with Tukey multiple comparison test). (J) Germline-specific loss of *mdy* in *bmm^1^* animals did not reduce GSC numbers, but the variance in GSC numbers was significantly rescued (*nos-GAL4*>*+*; *bmm^1^* vs *nos-GAL4*>*mdy RNAi*; *bmm^1^*: *p*=4.5 × 10^-5^; *+*>*mdy RNAi*; *bmm^1^* vs *nos-GAL4*>*mdy RNAi*; *bmm^1^*: *p*=0.0082 by F-test).

